# The vesicular trafficking system component MIN7 is required for minimizing *Fusarium graminearum* infection

**DOI:** 10.1101/2020.03.16.994095

**Authors:** Ana K. Machado Wood, Vinay Panwar, Mike Grimwade-Mann, Tom Ashfield, Kim E. Hammond-Kosack, Kostya Kanyuka

## Abstract

Plants have developed intricate defense mechanisms, referred to as innate immunity, to defend themselves against a wide range of pathogens. Plants often respond rapidly to pathogen attack by the synthesis and delivery of various antimicrobial compounds, proteins and small RNA in membrane vesicles to the primary infection sites. Much of the evidence regarding the importance of vesicular trafficking in plant-pathogen interactions comes from the studies involving model plants whereas this process is relatively understudied in crop plants. Here we assessed whether the vesicular trafficking system components previously implicated in immunity in *Arabidopsis thaliana* play a role in the interaction with *Fusarium graminearum*, a fungal pathogen notoriously famous for its ability to cause Fusarium head blight (FHB) disease in wheat. Among the analyzed vesicular trafficking mutants, two independent T-DNA insertion mutants in the *AtMin7* gene displayed a markedly enhanced susceptibility to *F. graminearum*. Earlier studies identified this gene, encoding an ARF-GEF protein, as a target for the HopM1 effector of the bacterial pathogen *Pseudomonas syringae* pv. *tomato*, which destabilizes AtMIN7 leading to its degradation and weakening host defenses. To test whether this key vesicular trafficking component may also contribute to defense in crop plants, we identified the candidate *TaMin7* genes in wheat and knocked-down their expression through Virus induced gene silencing. Wheat plants in which *TaMIN7* were silenced displayed significantly more FHB disease. This suggests that disruption of MIN7 function in both model and crop plants compromises the trafficking of innate immunity signals or products resulting in hyper-susceptibility to various pathogens.

**One sentence summary:** Disruption of an ARF-GEF protein encoding gene *AtMin7* in *Arabidopsis thaliana* and silencing of the orthologous gene in wheat result in hyper susceptibility to the fungal pathogen *Fusarium graminearum*.

## INTRODUCTION

Being sessile in nature, plants are frequently exposed to various environmental stresses including pathogens and yet more often than not plants appear healthy or show only weak or mild disease symptoms. To maintain this healthy status, plants have evolved an elaborate and tightly regulated innate immune system that allows them to fend off pathogens or restrict pathogens invasion or slow down/minimize disease progression (Jones and Dangl, 2006).

Each plant cell is surrounded by the plasma membrane (PM) and a cell wall and contain a variety of membrane-enclosed organelles. Transport of various cargo molecules across different membranes and sorting these to the correct cellular compartments, i.e. the PM, cell wall and the extracellular space (apoplast) is a fundamental process, central for multiple plant cell functions. Regulation of multiple cellular responses by the membrane trafficking network during plant-microbe interactions is required to facilitate a coordinated defense response at sites of pathogen attack. These involve rapid remodeling the composition of the PM including accumulation, activation and re-localization of cell surface immune receptors, reinforcement of the cell wall, secretion of antimicrobial proteins, phenolics and reactive oxygen species. Whereas other molecules are trafficked from the vacuole and other cellular compartments to the apoplast and to the cell wall to orchestrate cytoplasm communication during immune signal transduction. This has been well documented, with *A. thaliana* and its vast available genetic resources being particularly well exploited for functional studies of vesicular trafficking components and pathways (Ben Khaled et al., 2015; Gu et al., 2017; Yun and Kwon, 2017; Ekanayake et al., 2019).

The vesicular trafficking system comprises two main pathways, secretory and endocytic (**Figure 1a**), with both implicated in effective immunity against pathogens. Thus, for example, a range of defense-related proteins, antimicrobial metabolites and compounds strengthening plant cell wall such as callose are secreted to the sites of pathogen invasion. Concomitantly cell-surface immune receptors are subjected to endocytosis which is necessary for initiation of signal transduction and regulation of receptor activity (e.g. through recycling or degradation in the vacuole). Secretory pathways transport newly synthesized proteins and other macromolecules, such as phospholipids, polysaccharides, and small RNA (collectively referred to as ‘cargo’) from the endoplasmic reticulum (ER) via the Golgi apparatus to the plasma membrane (PM) or the extracellular space. In the endocytic pathway, membranous vesicles internalized in the PM undergo homotypic fusion to form early/sorting endosomes. Internalized cargo could then be sorted and recycled back to the PM through the recycling endosome, sent to the *trans-*Golgi network (TGN) via retrograde trafficking mechanisms, or trafficked through the late endosome/multivesicular body (MVB) to the vacuole (**Figure 1a**).

**Figure 1.**
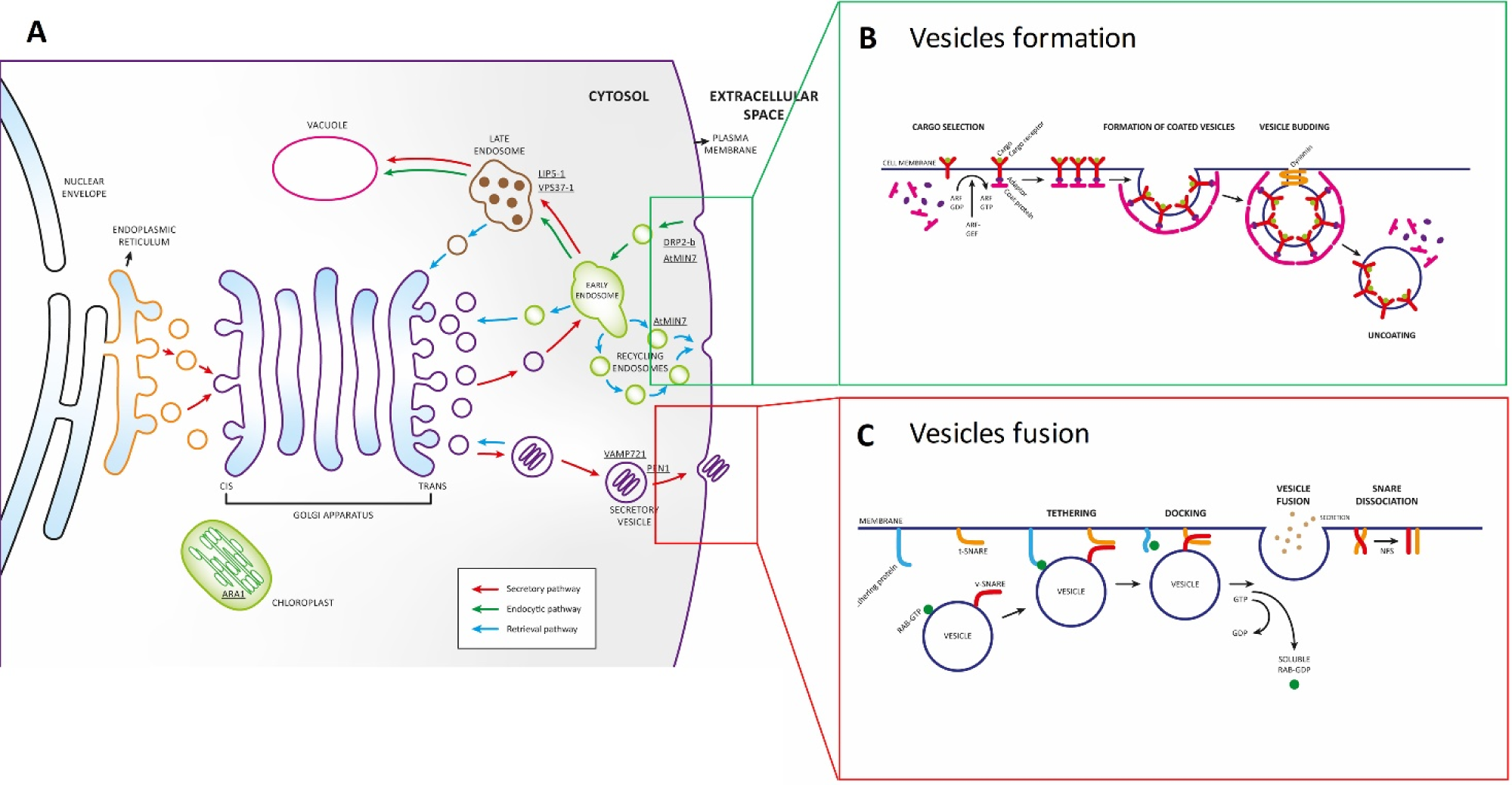
Schematic diagram of vesicular trafficking pathways in *Arabidopsis thaliana*. A, Major cellular vesicular trafficking pathways: secretory (red), endocytic (green) and retrieval (blue). B, vesicles formation and budding. C, Fusion of vesicles with the correct target membranes.

Vesicular trafficking requires the budding, transport and fusion of vesicles from the donor to the specific target membrane. Each of these processes is regulated by different components of the vesicular trafficking system. For instance, the bud formation requires small GTPases of ARF (ADP-ribosylation factor) or SAR1 (secretion-associated Ras-related protein) type as well as adaptor proteins that recognize and recruit specific cargo receptors. Phosphorylation and activation of ARF/SAR1 are regulated by guanine nucleotide exchange factor (GEF) proteins which stimulate the release of GDP to allow binding of GTP. The latter triggers a conformational change in ARF/SAR1 that allows their stable association with the membrane surface and recruitment of specific coat proteins (COP) initiating the budding process. Once the rounded vesicle shape is formed, the large Dynamin-related multidomain GTPases catalyze the membrane scission generating a transport vesicle. The coat components are rapidly lost shortly after the vesicle buds off (**Figure 1b**).

Fusion of vesicles to the correct target membranes is regulated by the small Rab GTPase proteins (**Figure 1c**). Different Rab proteins are associated with one or more membrane-enclosed organelles of the secretory pathway. Once in the GTP bound state, the Rab GTPase proteins bind to specific tethering factors in the target membrane to stablish the first connection between the membranes that are going to fuse. Next, the N-ethylmaleimide-sensitive factor adaptor protein receptor (SNARE)-domain containing proteins on both the vesicle and the targeting membrane dock to mediate the fusion of the two membranes. SNAREs are transmembrane proteins that exist in complementary sets. Those located in the vesicle are called v-SNAREs or R-SNAREs, and those located in the target membrane are known as t-SNAREs or Q-SNAREs. A trans-SNARE complex formed following binding of v-SNARE/R-SNARE to t-SNARE/Q-SNARE located on separate membranes catalyzes the membrane fusion (Collins et al., 2003; Yun and Kwon, 2017).

Much of the evidence regarding the role for vesicular trafficking in plant immunity comes from the studies involving model plants *Arabidopsis thaliana* and *Nicotiana benthamiana* and to a lesser extent crops such as barley (*Hordeum vulgare*) and a small number of their respective biotrophic or hemi-biotrophic bacterial (*Pseudomonas syringae*), oomycete (*Phytophthora infestans*) or the powdery mildew fungal pathogens (*Golovinomyces cichoracearum, Blumeria graminis*, and *Erysiphe pisi*) (Kwon et al., 2008a; Kwon et al., 2008b; Dodds and Rathjen, 2010; Bozkurt et al., 2011; Nomura et al., 2011; Ellinger et al., 2013). However, little is known about the role of plant vesicular trafficking in the interactions involving other pathogens and those which infect non-leaf tissue.

Fusarium Head Blight (FHB) disease caused by the ascomycete fungus *Fusarium graminearum* and related *Fusarium* species causes substantial yield losses and reduced grain quality and safety in a number of major cereal crops, such as wheat, barley, maize and oat, worldwide (Bottalico and Perrone, 2002). Moreover, under laboratory conditions *F. graminearum* is able to infect floral tissue of intact *A. thaliana* plants as well as detached leaves (Urban et al., 2003; Chen et al., 2006; Cuzick et al., 2008; Koch et al., 2013). For this plant species a large mutant collection is readily available.

The aim if this study was to assess whether knock-out mutations in the individual components of the vesicular trafficking system in *A. thaliana* previously implicated in plant immunity had any impact on the interaction with *F. graminearum*. Screening of the assembled mutants collection using a detached-leaf bioassay identified two independent T-DNA insertion mutants in the *AtMin7* gene, which encodes an ADP ribosylation factor (ARF) guanine nucleotide exchange factor (GEF) protein, that displayed striking hyper-susceptibility to *F. graminearum* strain PH-1 infection compared to the parental wild-type *A. thaliana* ecotype Col-0. Utilizing a recently released high-quality fully annotated wheat genome reference sequence assembly (International Wheat Genome Sequencing Consortium (IWGSC), 2018) and well established bioinformatics tools enabling identification of putative gene orthologs from different plant species (Adamski et al., 2019) we identified the three homoeologous *TaMin7* genes in hexaploid wheat (*Triticum aestivum*). Knock-down of these genes using Virus-induced gene silencing (VIGS) (Lee et al., 2012) significantly promoted FHB disease formation in this crop species.

## RESULTS

### Assembling a collection of *A. thaliana* mutants with defects in membrane trafficking

To investigate whether vesicular trafficking plays a role in a compatible interaction (i.e. disease) between an ascomycete fungus *F. graminearum* and its laboratory host *A. thaliana*, we assembled a collection of twelve mutants containing T-DNA insertions in nine immunity associated genes regulating different vesicular trafficking pathways (**Figure 1a**, and **Table 1**). Homozygous mutants were obtained from the Nottingham Arabidopsis Stock Centre (NASC, UK), and each mutant was verified by PCR amplification using gene-specific and T-DNA-specific primers as described in **Supplemental Methods** and **Supplemental Table S1**. No obvious developmental or growth defects were observed in any of the untreated mutant plants when compared with the corresponding wild type *A. thaliana* ecotype Col-0 plants grown under standard controlled environment conditions (data not shown).

**Table 1.** *Arabidopsis thaliana* mutants used in this study.

| Gene name | Gene ID | Gene description | Mutant name | Mutant accession No. | Reference | Evidence in Arabidopsis plants |
| --- | --- | --- | --- | --- | --- | --- |
| *Ara1/*<br>*AtRabA5E* | At1g05810 | Arabidopsis gene homologous to the Ras oncogene 1/ Rab GTPase-like A5A | *ara1* | SM_3.15870 | Karim et al., 2014 | Interacts with proteins involved in multiple stress responses |
| *Drp2b* | At1g59610 | Dynamin related protein 2B | *drp2b-1*<br>*drp2b-2*<br>*drp2b-3\** | SALK_003049<br>SALK_134887<br>SALK_124686 | Smith et al., 2014 | Functions as a novel component of flagellin-mediated defense responses against bacteria *P. syringae* |
| *Pen1/*<br>*Syp121* | At3g11820 | Penetration resistance 1/ Syntaxin of plants 121 | *pen1* | SALK_004484C | Johansson et al., 2014 | Absence of *pen1* is associated with increased tissue penetration by the fungal pathogen *Blumeria graminis* fsp. *hordei* (Bgh) |
| *Syp122* | At3g52400 | Syntaxin of plants 122 | *syp122-1\** | SALK_008617 | Zhang et al., 2007 | Acts together with SYP121 as a negative regulator of PCD, SA, JA and ET pathways |
| *AtMin7/*<br>*Ben1/*<br>*Big5* | At3g43300 | HopM1 interactor 7/ BFA-visualized endocytic trafficking defective 1/ Brefeldin A-inhibited guanine nucleotide-exchange protein 5 | *atmin7-1*<br>*atmin7-2* | SALK_012013<br>SALK_013761 | Nomura et al., 2006 | Encodes an immunity associated ARF-GEF protein family protein targeted by HopM1, conserved bacterial *P. syringae* virulence protein |
| *Vps37* | At3g53120 | Vacuolar protein sorting 37 | *vps37-1* | SALK_042859C | Spallek et al., 2013 | Co-localizes with FLS2 (FLAGELLIN SENSING 2 immune receptor) at endosomes and immunoprecipitates with this receptor upon flg22 elicitation |
| *Vps28-2* | At4g05000 | Vacuolar protein sorting 28 | *vps28-2\** | SALK_040274 | Spallek et al., 2013 | Critical for immunity against bacterial infection through a role in stomatal closure; regulates late FLS2 endosomal sorting |
| *Vamp721* | At1g04750 | Vesicle-associated<br>membrane protein 721 | *vamp721* | SALK_037273C | Kim et al.,<br>2014 | Forms complex with PEN1; mediates trafficking of the powdery mildew resistance protein RPW8.2 to the extrahaustorial membrane of haustorial complexes |
| *Lip5* | AT4G26750 | LYST-interacting protein<br>5 | *lip5-1* | SAIL_854_F08 | Wang et al., 2014 | Disruption of *LIP5* causes increased susceptibility to the bacterial pathogen *P. syringae* |

### AtMIN7, an ARF-GEF protein is required to minimize *F. graminearum* infections

To gain insight into whether mutations in any of the selected vesicular trafficking gene increase or decrease susceptibility to the virulent *F. graminearum* strain PH-1, young 5-weeks-old *A. thaliana* plants were point inoculated with *F. graminearum* conidial suspension supplemented with the mycotoxin deoxynivalenol (DON) in a detached leaf bioassay (Chen et al., 2006). At 6 days post inoculation (dpi), the inoculated leaves were photographed, and disease levels were quantified by measuring the proportion of lesioned/necrotic area compared to the total leaf area by analyzing the images using the LemnaGrid software module (LemnaTec GmbH, Aachen, Germany).

In the first two experiments, each involving a partially overlapping subset of the mutants, *F. graminearum* inoculated leaves of the two independent T-DNA insertion mutants in the *AtMin7* gene developed extensive necrotic lesions covering up to 75-100% of the total leaf area and showed almost complete loss of green photosynthetic tissue, while most of the remaining mutants displayed much milder disease symptoms with smaller lesions, the rest of the leaf remained unaffected and green (**Figure 2, Supplemental Figure S1**, and data not shown). Experiments involving nine out of twelve mutants, including *atmin7-1* and *atmin7-2*, and the wild-type *A. thaliana* Col-0 were replicated trice with similar results.

**Figure 2.**
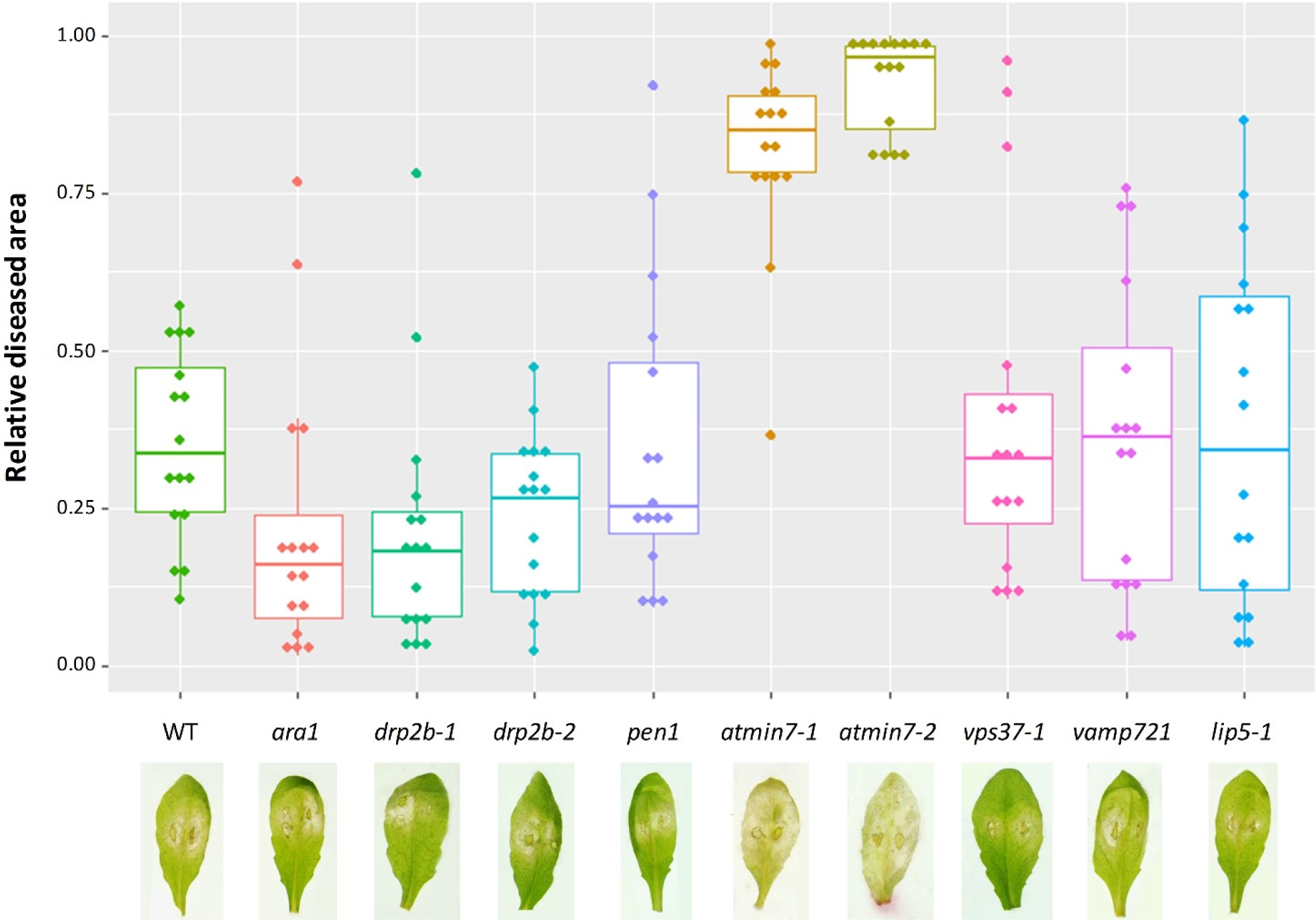
*F. graminearum* infection of different *Arabidopsis thaliana* mutants measured as the proportion of infected leaf area compared to the total leaf area. GLM analysis (*P* < 0.01). Data representing results from three different experiments with at least 30 leaves from 12 different plants from each mutant for each experiment.

*AtMin7* is a large gene of 5,857-nt containing 33 exons and is located on chromosome 3. The two loss of function mutants, *atmin7-1* and *atmin7-2*, studied here carry T-DNA insertions in the exon 1 and exon 18, respectively (Nomura et al., 2006). *AtMin7* is a member of small family comprising 8 genes encoding ARF-GEF proteins (**Supplemental Figure S2**), which play important roles in the budding of transport vesicles from the membranes (Steinmann et al., 1999; Mossessova et al., 2003). This vesicular trafficking component has been shown to contribute to resistance to the bacterial pathogen *Pseudomonas syringae* pv. *tomato* (*Pst*), possibly through regulating the trafficking of immunity-associated cargo molecules (Nomura et al., 2006; Nomura et al., 2011).

### Identification of candidate wheat *TaMin7* genes

To identify homologs of the *A. thaliana AtMin7* gene in wheat, a natural, crop host of *F. graminearum*, we used the BioMart data mining tool available through Ensembl Plants (Smedley et al., 2015). A total of 26 gene sequences were identified in the reference hexaploid wheat (*Triticum aestivum*) genome (AABBDD) by searching for genes encoding proteins containing the catalytic SEC7 domain (PF12783) characteristic of ARF-GEF proteins. We then aligned the proteins coded by the identified wheat genes with all eight members of the *A. thaliana* ARF-GEF family proteins and used the resulting multiple alignment for phylogenetic analysis. The constructed maximum likelihood phylogenetic tree revealed the three closely sequence related wheat proteins formed a distinct clade with AtMIN7, suggesting these sequences represent the wheat A-, B- and D-genome encoded orthologues of AtMIN7 (**Figure 3**).

**Figure 3.**
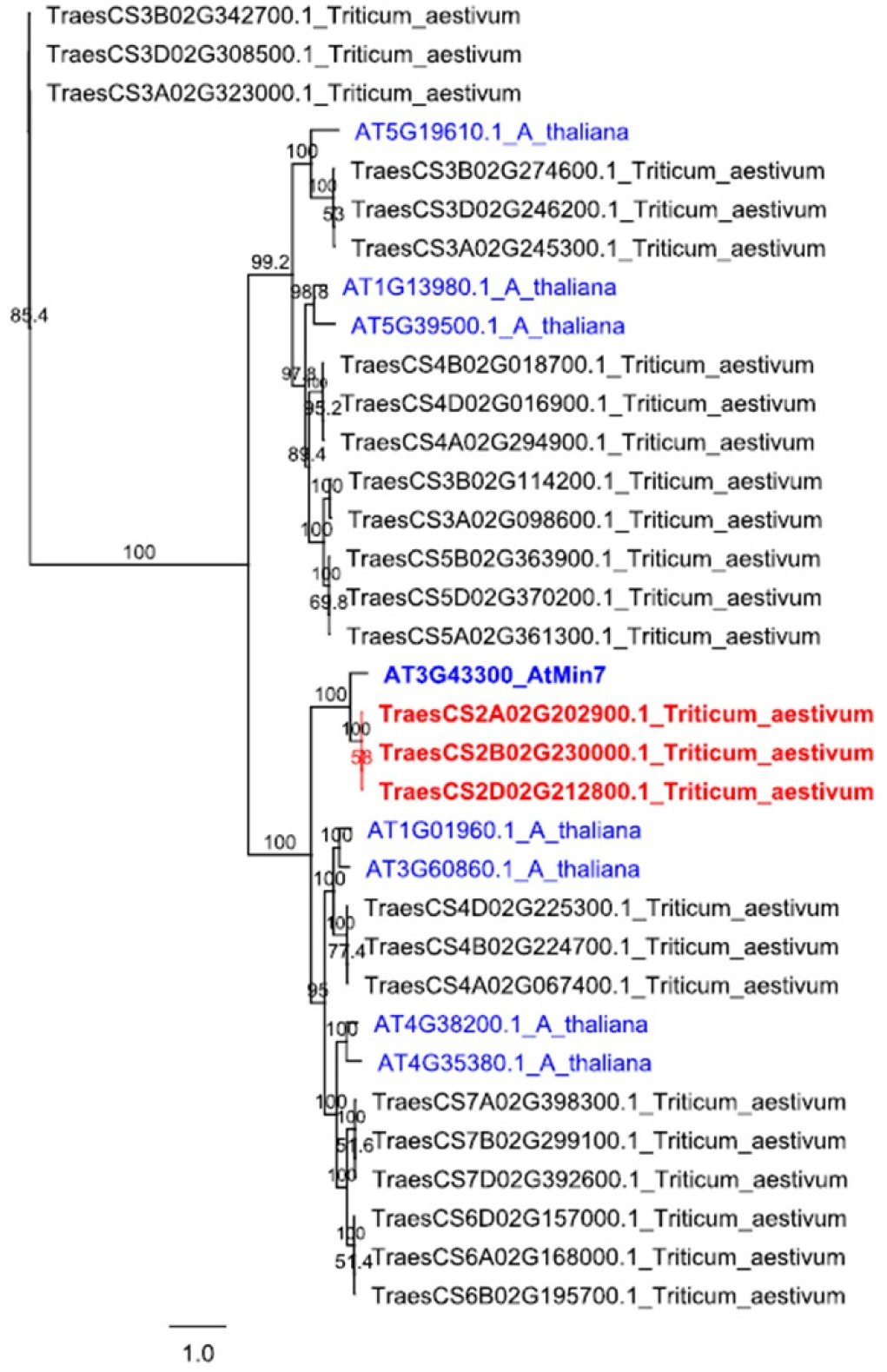
Maximum Likelihood phylogenetic tree displaying representative wheat sequence orthologues of *Arabidopsis thaliana ARF-GEF* genes (protein coding sequences). Node labels indicate percentage bootstrap support (500 replicates). The *A. thaliana ARF-GEF* gene names are shown in blue regular font with *AtMin7* accentuated in bold. Names of wheat genes most likely representing orthologs of *AtMin7* are shown in red bold font, while names of all other candidate wheat *ARF-GEF* genes are shown in regular black font.

### Silencing the candidate *TaMin7* genes in wheat spikes enhances susceptibility to *F. graminearum*

To explore the potential function for the wheat homologs of *AtMin7* in the *F. graminearum* - wheat interaction we tested the effect of silencing the three homoeologous wheat genes, *TraesCS2A02G202900, TraesCS2B02G230000* and *TraesCS2D02G212800*, using *Barley stripe mosaic virus-*mediated Virus induced gene silencing (VIGS) on the FHB disease development (Lee et al., 2012). A 209-nt fragment highly conserved between coding sequences of the three *TaMin7* homoeologs was selected as a target for VIGS using si-Fi21 software (Lück et al., 2019). This target fragment was predicted to generate a high number of silencing-effective siRNAs (*n* = 91), and a very low likelihood of off-target silencing.

An important factor for successful application of VIGS is the ability of the virus to infect and spread without causing any deleterious effect in the host plant. Therefore, the feasibility of BSMV-VIGS approach to induce systemic silencing in the spike tissue of wheat cv. Bobwhite susceptible to *F. graminearum* PH-1 (Brown et al., 2011) was first tested by visualizing the phenotype induced by silencing the *Magnesium-chelatase subunit H gene* (*TaChlH; TraesCS7A02G480700*) involved in chlorophyll biosynthesis, which is often used as a visible marker of VIGS (Yuan et al., 2011). The recombinant BSMV carrying a 250-nt fragment of *TaChlH* gene in antisense orientation and BSMV:*mcs4D* (harboring a 275-nt noncoding fragment corresponding to the multiple cloning site sequence of the vector pBluescript II KS) were inoculated onto the flag leaves of wheat plants at the early boot stage. At 13 days post inoculation (dpi) the plants infected with BSMV:*asTaChlH* developed yellow-orange coloration of the lemma and palea of spikes indicative of the loss of chlorophyll and successful silencing of the *TaChlH* gene, whereas the spikes of plants infected with the control construct BSMV:*mcs4D* showed typical mosaic symptoms (**Supplemental Figure S3**). No visible developmental abnormalities were observed in the wheat plants challenged with either of the two BSMV-VIGS constructs. Moreover, similar levels of FHB disease were observed on wheat plants pre-infected with BSMV and on virus-free plants challenged with *F. graminearum*, indicating that the susceptibility to the fungus was not compromised in the virus-infected wheat (**Figure 4**). These results suggest that BSMV-mediated VIGS can be used to silence genes in the reproductive tissue of wheat cv. Bobwhite and the approach appeared suitable for assessing the role of *TaMin7* during *F. graminearum* infection in wheat.

**Figure 4.**
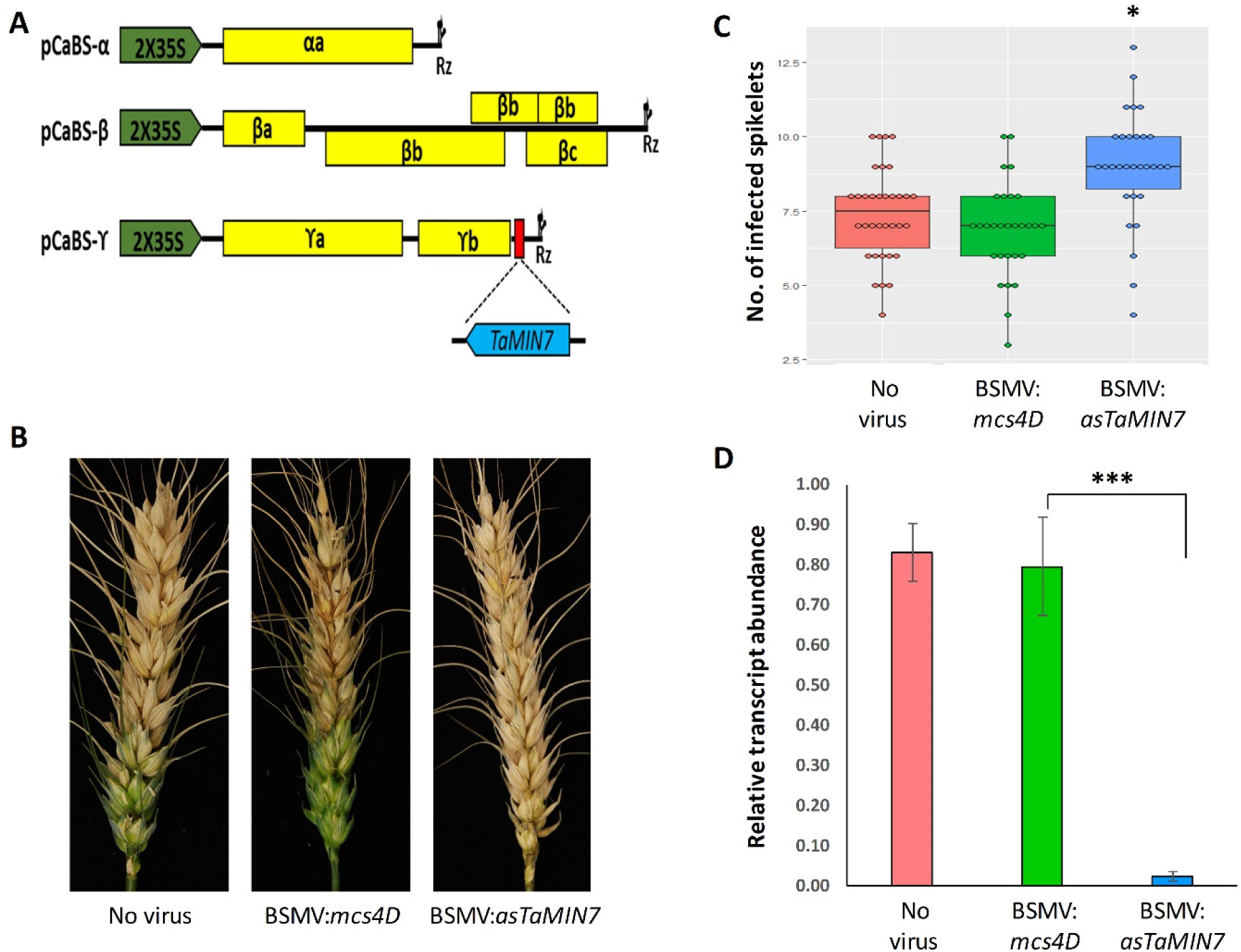
Effect of silencing *TaMin7* genes on the *Fusarium graminearum* infection in wheat. A, schematic representation of the *Barley stripe mosaic virus* (BSMV)-derived vectors for silencing *TaMin7* (adapted from Lee et al., 2012). cDNAs of the three BSMV genomic RNAs (α, β, and γ) each of which is required for full infection are cloned into a binary vector pCaBS under control of a double CaMV 35S promoter (2×35S) and flanked by a ribozyme sequence originating from TRSV satellite RNA (Rz), which allows cis-cleavage of transcribed RNA at the 3’ end of viral genomic RNA. The RNAγ genome is modified to include a small fragment of *TaMin7* protein-coding sequence in the antisense orientation immediately downstream of the BSMV *γb* cistron. Viral cistrons are shown as yellow rectangles. B, images of representative wheat spikes at 15 days post inoculation with *Fusarium graminearum* from virus-untreated (No virus) plants and those treated with the control (BSMV:*mcs4D*) and the VIGS construct for silencing TaMIN7 (BSMV:*asTaMin7*). C, *TaMin7* silenced wheat plants show increased susceptibility to *F. graminearum* infection as determined by counting the number of infected (bleached) spikelets per spike. Values represent mean of three independent experiments. D), qRT-PCR measurement of *TaMin7* transcripts abundance in wheat plants. RNA extracted prior to *F. graminearum* inoculation and the wheat *CDC48* (*Cell division control 48*) gene used as an internal reference. A single star (*) indicates *P* < 0.05 and three stars (***) indicate *P* < 0.001; *n* = number of plants.

The ability of BSMV:*asTaMin7* to induce silencing of the corresponding endogenous genes was confirmed using quantitative reverse transcription PCR (qRT-PCR). The *TaMin7* transcripts were decreased in abundance by 77 % in the spikes of the VIGS-treated plants compared to those inoculated with BSMV:*mcs4D*, and no significant difference in *TaMin7* expression levels was observed in the mock-versus BSMV:*mcs4D*-inoculated control plants. To analyze the effect of *TaMin7* silencing on *F. graminearum* infection, spikes of VIGS-treated plants showing typical virus-induced symptoms were point inoculated with the fungal conidia suspension and observed for disease symptoms for 15 days after fungal inoculation. Reduction of *TaMin7* mRNA expression levels in spikes of the BSMV:*asTaMin7* treated plants was associated with significantly enhanced susceptibility to *F. graminearum* (**Figure 4**). In contrast, no effect on FHB disease development was found in mock-inoculated and BSMV:*mcs4D*-inoculated control plants (**Figure 4**).

## DISCUSSION

Membrane trafficking plays an important role in plant-pathogen interactions and mutants with lesions in several different plant vesicular trafficking genes exhibiting compromised responses to bacterial or fungal pathogens in the model plant species *A. thaliana* have been identified in previous studies (Gu et al., 2017; Yun and Kwon, 2017; Ekanayake et al., 2019). Moreover, it appears that some pathogens evolved host cell translocated effector proteins that promote disease by interfering with plant membrane trafficking pathways (Ben Khaled et al., 2015). One of the vesicular trafficking genes whose role during plant disease has been previously well explored in the interactions between *A. thaliana* and the bacterial pathogen *Pst* is *AtMin7* (Nomura et al., 2006; Nomura et al., 2011). The AtMIN7 protein localizes to the TGN/early endosome (EE) and is involved in endocytic recycling of PM resident proteins but it has also been hypothesized to regulate secretion (Tanaka et al., 2009; LaMontagne and Heese, 2017). Mutants that lack this protein allow increased bacterial multiplication, possibly due to miss-regulation of membrane trafficking of the plant defense-related cargo (Nomura et al., 2006; Nomura et al., 2011). AtMIN7 has also been shown to contribute to the cytosol-initiated immune responses triggered by the *Pst* effectors such as AvrRpt2 and AvrPphB (Nomura et al., 2011) and to the apoplast-initiated immune responses through an unknown mechanism by preventing apoplast water soaking and therefore presumably restricting the flow of nutrients to the bacteria (Xin et al., 2016). To achieve successful disease, *Pst* secretes a conserved effector protein HopM1 that is translocated to the TGN/EE of its host during infection where it mediates destabilization of AtMIN7 followed by degradation via the 26S proteasome (Nomura et al., 2011). Here we demonstrated that AtMIN7 also contributes to defense against a fungal pathogen *F. graminearum* as the absence of this protein in *A. thaliana* resulted in a markedly enhanced disease (**Figure 2**). However, whereas *atmin7* mutants displayed increased susceptibility to the *Pst ΔCEL* mutant that lacks HopM1 along with several other conserved effectors including AvrE and only modest increase in susceptibility to the wild type *Pst* (Nomura et al., 2011), we show that these same *A. thaliana* mutants are clearly and unmistakably hyper-susceptible to the wild type strain of *F. graminearum* (**Figure 2**).

Disruption of *AtMin7* may compromise trafficking of specific molecules and cargo protein. These molecules could include plant defense-related proteins or secondary metabolites that help minimize *F. graminearum* infection. For example, callose (a (1,3)-β-glucan polymer) can act as a physical barrier reducing fungal penetration, and an increased callose deposition and the *F. graminearum* induced callose synthase activity in the infected spikelets was found to be correlated with increased disease resistance (Ribichich et al., 2000; Blümke et al., 2017). Identification of proteins and/or other molecules transported specifically by membrane vesicles regulated by AtMIN7 is challenging. But comparison of vesicles cargo between wild-type and *atmin7 A. thaliana* mutants during *F. graminearum* infection could provide valuable clues to the role of this gene during fungal disease establishment.

Although *A. thaliana* has proven to be a very useful model organism for unraveling the key mechanisms underlying plant-pathogen interactions, some findings cannot be translated directly to crop plants. Hence studies involving interactions between the pathogens and their natural host plants could provide more relevant information and facilitate exploration of new strategies for disease control in crops. Here we utilized a transient gene silencing approach (VIGS) to assess the role of *AtMin7* homologues in wheat, which is one of the most important staple food crops whose production is regularly threatened by fungal diseases including FHB. These results from the analysis of the wheat - *F. graminearum* pathosystem (**Figure 4**) are consistent with those from the study of the *A. thaliana* - *F. graminearum* pathosystem (**Figure 2**), and provide evidence that disruption of MIN7 function in both dicotyledonous and monocotyledonous hosts compromises the plant innate immunity resulting in more severe disease.

Whether *F. graminearum* utilizes similar mechanisms of suppressing the host immunity to promote disease as in the case of *Pst* which involve effector-mediated degradation of AtMIN7 remains to be determined. However, our preliminary data (not shown) indicates that *TaMin7* transcript levels may be reduced in the wheat spikes infected with *F. graminearum* compared to the mock-inoculated samples. If confirmed this would suggest a different mechanism operating in wheat as *Pst* infection of *A. thaliana* seem to induce AtMIN7 protein degradation while not affecting the *AtMin7* transcript levels (Nomura et al., 2006).

AtMIN7 was also shown to play a role in polar localization and dynamic repolarization of the PIN (PIN-formed) efflux carrier proteins enabling the directional transport of auxin in the tissues (Tanaka et al., 2013). Elevated levels of reactive oxygen species (ROS) induced during stress responses in *A. thaliana* affect AtMIN7-dependent PIN endocytic recycling resulting in increased accumulation of auxin in the affected tissues (Zwiewka et al., 2019). A recent study demonstrated that higher levels of auxin are accumulated during *F. graminearum* infection in a susceptible wheat cultivar Roblin compared to the moderately resistant cultivars Wuhan 1 and Nyubai, indicating that auxin may promote susceptibility to this fungal pathogen (Brauer et al., 2019). Moreover, analysis of the recently released transcriptome data indicated that wheat homologs of *AtMin7* and *PIN* genes are down-regulated during *F. graminearum* infection whereas numerous transcripts of genes belonging to auxin pathway are being up-regulated (Pan et al., 2018). The data from these previous studies together with findings from our study form a foundation to an alternative hypothesis regarding the mechanisms employed by *F. graminearum* for achieving successful infection. That is, it is conceivable that *F. graminearum* infection induces temporally coordinated waves of gene expression which regulate MIN7-dependent distribution and accumulation of auxin during infection. Further studies are necessary to confirm or refute this hypothesis.

This study also uncovered potential additional regulators of membrane trafficking contributing to host immunity such as SYP122. Significant increase (*P* < 0.05) in the size of lesions induced by *F. graminearum* in leaves of *syp122-1* compared to those in wild type *A. thaliana* Col-0 and other mutants (apart from *atmin7-1* and *atmin7-2*) was seen in one nonreplicated experiment (**Supplemental Figure S1**). This warrants further examination in a follow-on study, as well as analysis of response to *F. graminearum* in additional vesicular trafficking gene mutants that were not investigated in the current study and including those whose association with host immunity has not yet been demonstrated. It would also be interesting to investigate contribution of unconventional secretion pathways related to pathogen attack or stress conditions (Wang et al., 2018), which involve direct fusion of the central vacuole, multivesicular bodies or EXPO (a double-membrane exocyst-positive organelle) with the PM, to plant interactions with *F. graminearum*. Particularly interesting in this respect are the exosomes, small membranous vesicles that are thought to be released to the apoplast following fusion of MVBs with the PM (Rutter and Innes, 2017). These were reported to carry various defense related proteins and antimicrobial compounds, and small regulatory RNAs and appear to be able to cross the plant cell wall and deliver this defense-related cargo to the cells of invading fungal pathogens (Rutter and Innes, 2017; Cai et al., 2018).

Elucidating the mechanisms by which membrane trafficking regulates plant immune responses and enhanced understanding of the vesicular trafficking components and pathways manipulated by microbial pathogens to promote disease will provide fundamental knowledge for the development of novel methods of disease intervention.

## MATERIALS AND METHODS

### Plant material and growth conditions

The *A. thaliana* mutants (**Table 1**) and the corresponding wild-type parental ecotype Col-0 used in the study were obtained from the Nottingham Arabidopsis Stock Center. The corresponding mutations were verified by PCR using the primers listed in **Supplemental Table S1** and confirmed homozygous mutants selected for further study (**Supplemental Methods**). *A. thaliana* seeds were sown in Levington F2 + S compost (Everris Ltd.) and stratified in the dark for four days at 5°C before transferring to a controlled environment growth chamber operating at 20°C/17°C during day/night and a 16-h photoperiod (light intensity of approximately 100 µmol m^−2^ s^−1^).

The bread wheat (*Triticum aestivum*) cultivar (cv.) Bobwhite, was used in this study. The plants were grown in a controlled environment growth chamber with day/night temperatures of 23°C/20°C at around 65% relative humidity and a 16-h photoperiod with light intensity of approximately 180 µmol m^−2^ s^−1^.

### Fungal growth conditions, plants inoculation, and disease assessment

The *F. graminearum* strain PH-1 was used in this study. Routine culturing of the fungus, conidiospore induction, and preparation of conidial suspensions followed essentially the same procedures as described in Brown et al. (2011). Conidiospore suspensions harvested in sterile distilled water were adjusted to a concentration of 5 × 10^5^ or 1 × 10^5^ conidia/ml^−1^ for inoculation of *A. thaliana* or wheat, respectively.

Detached *A. thaliana* leaves were inoculated as described in Chen et al. (2006) with the following modification. Fully expanded rosette leaves were detached from the five-week-old plants using razor blades and placed adaxial surface facing upwards on the surface of 1% water agar in 10 × 10 cm square sterile Petri dishes, with 8 leaves per dish. Each leaf was then superficially wounded by gently puncturing over the mid rib with a glass Pasteur pipette and a 5 µL droplet of *F. graminearum* conidiospore suspension supplemented with 20 µM deoxynivalenol (DON) was deposited on the fresh wound. Mock inoculation was carried out using a 5 µL droplet of sterile distilled water supplemented with 20 µM DON. After inoculation, the plates were transferred to the controlled environment growth chamber operating at 20°C/17°C during day/night and 16-h photoperiod but kept in the dark for the first three days following which they were incubated under low light (light intensity of 40 µmol m^−2^ s^−1^) for further four days before the disease assessment took place.

Color (RGB) photographs were taken at six days after inoculation using a Nikon (D90) camera and backlighting to ensure consistent illumination. Image analysis to quantify the diseased areas was conducted using the LemnaTec LemnaGrid software module (LemnaTec GmbH, Aachen, Germany). Leaf areas were segmented using a combination of a colour-based classification and thresholding after converting the images to grayscale. Filters were applied to remove misclassified pixels and to fill in gaps. Finally, a customized colour-based classification was applied to score leaf-area pixels as belonging to diseased or healthy tissue.

Intact spikes of adult wheat cv. Bobwhite plants were point inoculated at the first signs of anther extrusion by depositing 5 μL of conidial suspension in the floral cavity between the palea and lemma of the outer two florets located in the upper one third of the spike as previously described (Brown et al., 2011). Control plants were inoculated with sterile water only. Inoculated plants were incubated in a humid chamber for 48 hr of which the first 24 h were in darkness. The inoculated plants were then kept in a controlled environment growth chamber at approximately 65% humidity, and the progression of the disease was visually monitored every 3 days and the number of bleached spikelets below the inoculated spikelet on each spike was recorded (Urban et al., 2003).

### Identification of putative wheat *Min7* genes

Protein domain analysis of predicted *ARF-GEF* genes in the hexaploid wheat (*T. aestivum*) was carried out using BioMart tool in Ensembl. Initially we searched for proteins that contained the Sec7_N domain (Guanine nucleotide exchange factor in Golgi transport N-terminal domain; PF12783). The wheat genome assembly used for this analysis was the IWGSCrefseq1 (International Wheat Genome Sequencing Consortium (IWGSC), 2018). Coding sequences of eight previously identified genes comprising the *ARF-GEF* gene family in *A. thaliana* (Vernoud et al., 2003) were also extracted using BioMart tool in Ensembl. Multiple protein sequences alignment was carried out in ClustalW, linked to Geneious 10. For phylogenetic reconstruction, the TVM+I+G nucleotide substitution model was selected by AIC in jModeltest 2.1.10 (Posada, 2008; Darriba et al., 2012). The Maximum Likelihood phylogeny was reconstructed using PhyML (Guindon and Gascuel, 2003), with the substitution model selected in jModeltest; starting tree with optimized topology, length and rate parameters; topology searching by the best of NNI and SPR; and 500 bootstraps.

### Vector construction for VIGS and fungal inoculations of wheat plants

The total RNA extracted from healthy wheat cv. Bobwhite leaf tissue was used as a template in an RT-PCR for amplification of 209-bp *TaMin7* gene fragment for BSMV-VIGS vector construction. The primers TaMin7-2A-seg1-R (5’-AACCACCACCACCGTAAAAGGGTCGCCTCGTCAAT-3’) and TaMin7-2A-seg1-F (5’-AAGGAAGTTTAATGTTGCAAGCAAAGGCCATC-3’) were designed to clone the *TaMin7* gene segment in an antisense orientation into BSMV RNAγ vector pCa-γbLIC using a ligation independent cloning (Yuan et al., 2011). VIGS vector for silencing the control, *TaChlH* gene was kindly provided by Dawei Li (China Agricultural University, Beijing, China). To prepare the virus inoculum for wheat inoculation the BSMVα, BSMVβ and the recombinant BSMVγ derivatives containing the *TaMin7* and *TaChlH* inserts were transformed into the *Agrobacterium tumefaciens* strain GV3101 (pMP90). Agroinfiltration of *Nicotiana benthamiana* leaves was carried out as previously described (Lee et al., 2015). The infiltrated *N. benthamiana* leaves were harvested 5-7 days post agroinfiltration, homogenized in 10 mM Na-phosphate buffer (pH 6.8) containing 1% Celite 545 AW (Sigma-Aldrich, UK), and the sap was mechanically inoculated onto wheat leaves just prior to appearance of a flag leaf. Fungal inoculation of wheat plants was carried out using *F. graminearum* wild type strain PH-1 as described in Brown et al. (2011). Briefly, 5 µL of conidial suspension was inoculated to the outer two florets between the lemma and palea in the upper part of the spike. Control plants were inoculated with sterile water only.

### Quantitative real time PCR (QPCR)

Total RNA was isolated from spike tissue of wheat plants infected with BSMV:*asTaMin7*, BSMV:*mcs4D* and mock-inoculated plants prior to *F. graminearum* inoculations using the TRIzol reagent (Invitrogen, Carlsbad, CA, USA) following the manufacturer instructions. To remove any traces of gDNA contamination, RNA samples were treated with TURBO DNaseI (Invitrogen) using methods as described by the manufacturer. The first strand cDNA was synthesized from 1 µg of total RNA in a total volume of 20 µL using the SuperScript IV Reverse Transcriptase (Invitrogen) and oligo (dT)_18_ primers according to manufacturer’s instructions. *TaMin7-*specifc primers for qPCR, TaMIN7-Pair1F-QPCR (5’-ATCTTGCGGCAAAAACCAGT-3’) and TaMIN7-Pair1R-QPCR (5’-ACCTGCTGAGCCACATGAAA-3’), were designed using Primer3. Expression of *TaMin7* in silenced and mock-inoculated leaves was normalized to the housekeeping *Cell division control 48* gene (*CDC48*) essentially as previously described (Lee et al., 2014). A no template control was included in all qPCR experiments.

### Statistical Analyses

For *A. thaliana* leaf inoculations assays, disease was quantified by expressing the diseased leaf area relative to the total leaf area. Mean disease levels for each genotype were compared using a multi-stratum analysis of variance (ANOVA). Independent lines were compared with the wild type using a Dunnett’s test at the 5% (*P* < 0.05) level of significance using wild-type *A. thaliana* Col-0 as the control test. GenStat (release 20.1, 2019, VSN International Ltd.) was used for the statistical analyses.

## ACKNOWLEDGMENTS

We thank Dawei Li for providing the BSMV:*asTaChlH* construct, and Amy Dodd for her expert assistance with the illustrations.

## SUPPLEMENTAL DATA

**Supplemental Methods.** PCR-based confirmation of specific mutations in the T-DNA insertion mutant *Arabidopsis thaliana* plants obtained from the seed stock center.

**Supplemental Table S1.** Primers used in this study for genotyping *Arabidopsis thaliana* mutants.

**Supplemental Figure S1.** *Fusarium graminearum* infection of additional different *Arabidopsis thaliana* mutants measured as proportion of infected leaf area compared to the total leaf area in a single nonreplicated experiment. Significant differences between wild-type *A. thaliana* ecotype Col-0 plants and vesicle trafficking mutants indicated by letter according Tukey’s test subsequent to ANOVA (*P* < 0.05).

**Supplemental Figure S2.** Maximum Likelihood phylogenetic tree indicating the relationship among *Arabidopsis thaliana ARF-GEF* genes (coding sequences only). Protein sequences were aligned using the ClustalW in the Geneious 10 software. Node labels indicate percentage bootstrap support (500 replicates).

**Supplemental Figure S3.** Silencing of *TaChlH* (*Mg-chelatase subunit H*) gene in wheat spikes. *TaChlH* silencing phenotype could be identified approximately two weeks after inoculation of flag leaves with BSMV:*asTaChlH* whereas those inoculated with a negative control BSMV:*mcs4D* construct and mock-inoculated plants showed no spike yellowing phenotype.

## Parsed Citations

Adamski NM, Borrill P, Brinton J, Harrington S, Marchal C, Bentley AR, Bovill WD, Cattivelli L, Cockram J, Contreras-Moreira B, et al (2019) Aroadmap for gene functional characterisation in wheat. PeerJ Prepr 7: e26877v2

Ben Khaled S, Postma J, Robatzek S (2015) A moving view: subcellular trafficking processes in pattern recognition receptor-triggered plant immunity. Annu Rev Phytopathol 53: 379–402

Blümke A, Falter C, Herrfurth C, Sode B, Bode R, Schäfer W, Feussner I, Voigt CA (2017) Secreted fungal effector lipase releases free fatty acids to inhibit innate immunity-related callose formation during wheat head infection. Plant Physiol 165: 346–358

Bozkurt TO, Schornack S, Win J, Shindo T, Ilyas M, Oliva R, Cano LM, Jones AME, Huitema E, van der Hoorn RAL, et al (2011) Phytophthora infestans effector AVRblb2 prevents secretion of a plant immune protease at the haustorial interface. Proc Natl Acad Sci U S A 108: 20832–20837

Brauer EK, Rocheleau H, Balcerzak M, Pan Y, Fauteux F, Liu Z, Wang L, Zheng W, Ouellet T (2019) Transcriptional and hormonal profiling of Fusarium graminearum-infected wheat reveals an association between auxin and susceptibility. Physiol Mol Plant Pathol 107: 33–39

Brown NA, Bass C, Baldwin TK, Chen H, Massot F, Carion PWC, Urban M, van de Meene AML, Hammond-Kosack KE (2011) Characterisation of the Fusarium graminearum-wheat floral interaction. J Pathog 2011: 626345

Cai Q, Qiao L, Wang M, He B, Lin F-M, Palmquist J, Huang S-D, Jin H (2018) Plants send small RNAs in extracellular vesicles to fungal pathogen to silence virulence genes. Science 360: 1126–1129

Chen X, Steed A, Harden C, Nicholson P (2006) Characterization of Arabidopsis thaliana-Fusarium graminearum interactions and identification of variation in resistance among ecotypes. Mol Plant Pathol 7: 391–403

Collins NC, Thordal-Christensen H, Lipka V, Bau S, Kombrink E, Qiu J-L, Hückelhoven R, Stein M, Freialdenhoven A, Somerville SC, et al (2003) SNARE-protein-mediated disease resistance at the plant cell wall. Nature 425: 973–977

Cuzick A, Urban M, Hammond-Kosack K (2008) Fusarium graminearum gene deletion mutants map1 and tri5 reveal similarities and differences in the pathogenicity requirements to cause disease on Arabidopsis and wheat floral tissue. New Phytol 177: 990–1000

Darriba D, Taboada GL, Doallo R, Posada D (2012) jModelTest 2: more models, new heuristics and parallel computing. Nat Methods 9: 772

Dodds PN, Rathjen JP (2010) Plant immunity: towards an integrated view of plant-pathogen interactions. Nat Rev Genet 11: 539–548

Ekanayake G, LaMontagne ED, Heese A (2019) Never walk alone: clathrin-coated vesicle (CCV) components in plant immunity. Annu Rev Phytopathol 57: 387–409

Ellinger D, Naumann M, Falter C, Zwikowics C, Jamrow T, Manisseri C, Somerville SC, Voigt CA (2013) Elevated early callose deposition results in complete penetration resistance to powdery mildew in Arabidopsis. Plant Physiol 161: 1433–1444

Gu Y, Zavaliev R, Dong X (2017) Membrane trafficking in plant immunity. Mol Plant 10: 1026–1034

Guindon S, Gascuel O (2003) Asimple, fast, and accurate algorithm to estimate large phylogenies by maximum likelihood. Syst Biol 52: 696–704

International Wheat Genome Sequencing Consortium (IWGSC) (2018) Shifting the limits in wheat research and breeding using a fully annotated reference genome. Science 361: eaar7191

Johansson ON, Fantozzi E, Fahlberg P, Nilsson AK, Buhot N, Tör M, Andersson MX (2014) Role of the penetration-resistance genes PEN1, PEN2 and PEN3 in the hypersensitive response and race-specific resistance in Arabidopsis thaliana. Plant J 79: 466–476

Jones JDG, Dangl JL (2006) The plant immune system. Nature 444: 323–329

Karim S, Alezzawi M, Garcia-Petit C, Solymosi K, Khan NZ, Lindquist E, Dahl P, Hohmann S, Aronsson H (2014) A novel chloroplast localized Rab GTPase protein CPRabA5e is involved in stress, development, thylakoid biogenesis and vesicle transport in Arabidopsis. Plant Mol Biol 84: 675–692.

Kim H, O’Connell R, Maekawa-Yoshikawa M, Uemura T, Neumann U, Schulze-Lefert P (2014) The powdery mildew resistance protein RPW8.2 is carried on VAMP721/722 vesicles to the extrahaustorial membrane of haustorial complexes. Plant J 79: 835–847

Koch A, Kumar N, Weber L, Keller H, Imani J, Kogel K-H (2013) Host-induced gene silencing of cytochrome P450 lanosterol C14α-demethylase-encoding genes confers strong resistance to Fusarium species. Proc Natl Acad Sci U S A 110: 19324–19329

Kwon C, Bednarek P, Schulze-Lefert P (2008a) Secretory pathways in plant immune responses. Plant Physiol 147: 1575–1583

Kwon C, Neu C, Pajonk S, Yun HS, Lipka U, Humphry M, Bau S, Straus M, Kwaaitaal M, Rampelt H, et al (2008b) Co-option of a default secretory pathway for plant immune responses. Nature 451: 835–840

LaMontagne ED, Heese A (2017) Trans-Golgi network/early endosome: a central sorting station for cargo proteins in plant immunity. Curr Opin Plant Biol 40: 114–121

Lee W-S, Hammond-Kosack KE, Kanyuka K (2012) Barley stripe mosaic virus-mediated tools for investigating gene function in cereal plants and their pathogens: Virus-induced gene silencing, Host-mediated gene silencing, and Virus-mediated overexpression of heterologous protein. Plant Physiol 160: 582–590

Lee W-S, Rudd JJ, Hammond-Kosack KE, Kanyuka K (2014) Mycosphaerella graminicola LysM effector-mediated stealth pathogenesis subverts recognition through both CERK1 and CEBiP homologues in wheat. Mol Plant Microbe Interact 27: 236–243

Lee W-S, Rudd JJ, Kanyuka K (2015) Virus induced gene silencing (VIGS) for functional analysis of wheat genes involved in Zymoseptoria tritici susceptibility and resistance. Fungal Genet Biol 79: 84–88

Lück S, Kreszies T, Strickert M, Schweizer P, Kuhlmann M, Douchkov D (2019) siRNA-Finder (si-Fi) software for RNAi-target design and off-target prediction. Front Plant Sci. 10: 1023.

Mossessova E, Corpina RA, Goldberg J (2003) Crystal structure of ARF1*Sec7 complexed with Brefeldin A and its implications for the guanine nucleotide exchange mechanism. Mol Cell 12: 1403–1411

Nomura K, Debroy S, Lee YH, Pumplin N, Jones J, He SY (2006) A bacterial virulence protein suppresses host innate immunity to cause plant disease. Science 313: 220–223

Nomura K, Mecey C, Lee Y-N, Imboden LA, Chang JH, He SY (2011) Effector-triggered immunity blocks pathogen degradation of an immunity-associated vesicle traffic regulator in Arabidopsis. Proc Natl Acad Sci U S A 108: 10774–10779

Pan Y, Liu Z, Rocheleau H, Fauteux F, Wang Y, McCartney C, Ouellet T (2018) Transcriptome dynamics associated with resistance and susceptibility against Fusarium Head Blight in four wheat genotypes. BMC Genomics 19: 642

Posada D (2008) jModelTest: phylogenetic model averaging. Mol Biol Evol 25: 1253–1256

Ribichich KF, Lopez SE, Vegetti AC (2000) Histopathological spikelet changes produced by Fusarium graminearum in susceptible and resistant wheat cultivars. Plant Dis 84: 794–802

Rutter BD, Innes RW (2017) Extracellular vesicles isolated from the leaf apoplast carry stress-response proteins. Plant Physiol 173: 728–741

Smedley D, Haider S, Durinck S, Pandini L, Provero P, Allen J, Arnaiz O, Awedh MH, Baldock R, Barbiera G, et al (2015) The BioMart community portal: an innovative alternative to large, centralized data repositories. Nucleic Acids Res 43: W589–W598

Smith JM, Leslie ME, Robinson SJ, Korasick DA, Zhang T, Backues SK, Cornish PV, Koo AJ, Bednarek SY, Heese A (2014) Loss of Arabidopsis thaliana Dynamin-Related Protein 2B reveals separation of innate immune signaling pathways. PLoS Pathog 10: e1004578

Spallek T, Beck M, Ben Khaled S, Salomon S, Bourdais G, Schellmann S, Robatzek S (2013) ESCRT-I mediates FLS2 endosomal sorting and plant immunity. PLoS Genet 9: e1004035

Steinmann T, Geldner N, Grebe M, Mangold S, Jackson CL, Paris S, Gälweiler L, Palme K, Jürgens G (1999) Coordinated polar localization of auxin efflux carrier PIN1 by GNOM ARF GEF. Science 286: 316–318

Tanaka H, Kitakura S, Rakusová H, Uemura T, Feraru MI, De Rycke R, Robert S, Kakimoto T, Friml J (2013) Cell polarity and patterning by PIN trafficking through early endosomal compartments in Arabidopsis thaliana. PLoS Genet 9: e1003540

Tanaka H, Kitakura S, De Rycke R, De Groodt R, Friml J (2009) Fluorescence imaging-based screen identifies ARF GEF component of early endosomal trafficking. Curr Biol 19: 391–397

Urban M, Mott E, Farley T, Hammond-Kosack K (2003) The Fusarium graminearum MAP1 gene is essential for pathogenicity and development of perithecia. Mol Plant Pathol 4: 347–359

Vernoud V, Horton AC, Yang Z, Nielsen E (2003) Analysis of the small GTPase gene superfamily of Arabidopsis. Plant Physiol 131: 1191–1208

Wang F, Shang Y, Fan B, Yu JQ, Chen Z (2014) Arabidopsis LIP5, a positive regulator of multivesicular body biogenesis, is a critical target of pathogen-responsive MAPK cascade in plant basal defense. PLoS Pathog 10: e1004243

Wang X, Chung KP, Lin W, Jiang L (2018) Protein secretion in plants: conventional and unconventional pathways and new techniques. J Exp Bot 69: 21–37

Xin X-F, Nomura K, Aung K, Velásquez AC, Yao J, Boutrot F, Chang JH, Zipfel C, He SY (2016) Bacteria establish an aqueous living space in plants crucial for virulence. Nature 539: 524–529

Yuan C, Li C, Yan L, Jackson AO, Liu Z, Han C, Yu J, Li D (2011) A high throughput Barley stripe mosaic virus vector for virus induced gene silencing in monocots and dicots. PLoS One 6: e26468

Yun HS, Kwon C (2017) Vesicle trafficking in plant immunity. Curr Opin Plant Biol 40: 34–42

Zhang Z, Feechan A, Pedersen C, Newman MA, Qiu JL, Olesen KL, Thordal-Christensen H (2007) A SNARE-protein has opposing functions in penetration resistance and defence signalling pathways. Plant J 49: 302–312

Zwiewka M, Bielach A, Tamizhselvan P, Madhavan S, Ryad EE, Tan S, Hrtyan MN, Dobrev P, Vankovï R, Friml J, et al (2019) Root adaptation to H2O2-induced oxidative stress by ARF-GEF BEN1- and cytoskeleton-mediated PIN2 trafficking. Plant Cell Physiol 60: 255–273

